# Higher frequency of extra-pair offspring in urban than forest broods of great tits (*Parus major*)

**DOI:** 10.1101/526194

**Authors:** Ivett Pipoly, Krisztián Szabó, Veronika Bókony, Bálint Preiszner, Gábor Seress, Ernő Vincze, Julia Schroeder, András Liker

## Abstract

Urbanization increasingly changes the ecological conditions for wild animal populations, influencing their demography, reproduction, and behaviour. While studies on the ecological consequences of urbanization frequently document a reduced number and poorer body condition of offspring in urban than in non-urban bird populations, consequences for other components of reproduction are rarely investigated. Mating with partners outside the social pair-bond is widespread in birds, and although theory predicts that the occurrence of extra-pair fertilizations (EPF) may be sensitive to the altered ecological conditions of cities, the effect of urbanization on EPF is poorly known. Here we used data from two urban and two forest populations collected over three years to test whether the frequency of extra-pair offspring (EPO) in great tit broods differed between the habitats. We found that significantly more broods contained EPO in urban habitats (48.9 %) than in forests (24.4 %). In broods with EPO, the number and proportion of EPO was similar in urban and forest broods. These results suggest that females that live in urban habitats are more likely to engage in EPF than those living in forests. Urban environments may either 1) provide more spatiotemporal opportunities to EPF because of higher breeding density and lower or more constant caterpillar supply in cities compared to natural habitats, or 2) enhance the benefits of EPF via increased fertility or due to disrupted quality signals caused by anthropogenic pollution. In addition, 3) females with higher propensity to engage in EPF may more likely settle in urban habitats.

## Introduction

Urban animals often differ in their behaviour, life history, demographics and fitness from conspecifics living in natural habitats (Gil & Brumm 2013; Rodewald & Gehrt 2014; Seress & Liker 2015). In birds, many species successfully colonized urban areas worldwide, and urban individuals have to cope with several anthropogenic environmental changes such as noise, light and chemical pollution (Seress & Liker 2015), habitat fragmentation (Crooks, Suarez & Bolger 2004), and ecological challenges such as higher population densities (Moller *et al*. 2012) and lower availability of natural food (Seress et al. 2018). These differences between urban and natural habitats may alter the costs and benefits of birds’ reproductive decisions thereby affecting their behaviour. In line with this, urban birds typically show altered reproductive biology including advanced laying dates, reduced brood sizes, higher nest-failure rates and smaller nestlings (Bailly *et al*. 2016; Seress *et al*. 2018) compared to their non-urban conspecifics.

An alternative reproductive strategy widespread among pair-bonding species is the pursuit of extra-pair fertilizations (EPF), which are found in approximately 90% of socially monogamous bird species (Griffith, Owens & Thuman 2002). EPF can increase the number of offspring for males, and it may grant genetic benefits to females by improving fertilization success and/or offspring quality (Griffith *et al*. 2002). These adaptive functions of EPF have been supported by empirical data, although not unequivocally (Hsu 2014; Arct, Drobniak & Cichoń 2015). The frequency of EPF can also be influenced by the spatiotemporal distribution of mating opportunities (Schlicht, Valcu & Kempenaers 2014; García-Navas *et al*. 2015) and physical environmental factors such as night lighting and anthropogenic noise (Kempenaers *et al*. 2010; Halfwerk *et al*. 2011). Many of these potential drivers of extra-pair mating behaviour may be affected by the altered ecological conditions of urban habitats. However, very few studies have compared EPF between birds breeding in urban and natural habitats (Moore *et al*. 2012; Rodriguez-Martínez *et al*. 2014, Bonderud *et al*. 2018), and it remains unclear whether and how extra-pair behaviour varies with habitat urbanization.

The aim of our study was to test whether the frequency of extra-pair offspring (EPO; i.e. offspring that are not genetically related to the social partner of the female) within and across broods differs between urban and non-urban great tits (*Parus major*), a common and successfully urbanized passerine species with relatively high EPF rates (García-Navas *et al*. 2015). To test this we used a data set from two urban and two forest populations from three consecutive breeding seasons.

## Methods

We studied great tits breeding in nestboxes at two urban and two forest sites in Hungary from 2012 to 2014. We recorded the number of eggs and nestlings in the nestboxes every 3-4 days from March to the end of June. We captured parent birds and took their nestlings before fledging to collect blood samples for genotyping. In 2013 and 2014 we also collected unhatched eggs and tissue samples from nestlings that were found dead before blood sampling. For further details, see Supplementary Information: Field methods.

We selected 86 first annual broods of marked parents and conducted multi-locus genotyping on the whole families by amplifying 5 microsatellite loci with tri- and tetra-nucleotide repeats (Table S1) using multiplex PCR reactions. Altogether 159 parents (80 males and 79 females) and 851 offspring were genotyped. In a subset of samples (n=23 individuals out of total 1010) with ambiguous results based on the 5 loci, we used 3 additional loci (Table S1). Fluorescent PCR products were scanned by capillary electrophoresis, and alleles were identified and scored by two independent researchers who were blind to the identity of birds. Our marker set proved reliable and efficient for identifying within-pair offspring (WPO) and EPO (see details in Supplementary Information: Genotyping). We identified an offspring as EPO if it mismatched the alleles of the social father on at least two loci but it had no mismatch with the maternal alleles.

We tested the difference in EPO frequencies between urban and forest broods by Generalised Estimation Equations (GEE) models to accommodate the non-normal error distributions, the few non-independent broods as the majority of broods were independent but a few had one or two parents in common (Zuur *et al*. 2009). We investigated three response variables: EPO occurrence (EPO present / absent in the brood), the number of EPO per brood, and the proportion of EPO within the brood, calculated as EPO / (EPO + WPO). The model for EPO occurrence contained all 86 broods, while the models for EPO number and EPO proportion contained only those 32 broods where at least one EPO occurred. We chose this subset in the latter models to avoid zero inflation, and to explicitly address the question whether urban and forest parents that engage in EPF differ in their allocation into extra-pair offspring. The predictor variables were study site (4 sites) and year (3 years) in all models. We calculated a linear contrast from each GEE model’s estimates to statistically compare the two habitat types (i.e. two urban sites versus two forest sites). For further details, see Supplementary Information: Statistical analyses.

## Results

There were more broods containing EPO in urban than forest habitat (Figure 1, Table 1) in all three study years (urban vs. forest 2012: 40.00 vs. 33.33 %; 2013: 39.13 vs. 6.67 %; 2014: 64.71 vs. 35.29 %). When we controlled for non-independence of parents and differences among years in a GEE model (Table S2), EPO occurrence was significantly higher in urban than forest broods (odds ratio calculated from urban-forest linear contrast = 5.77, 95 % confidence interval (CI): 1.72 – 19.4, p = 0.005).

**Figure 1.**
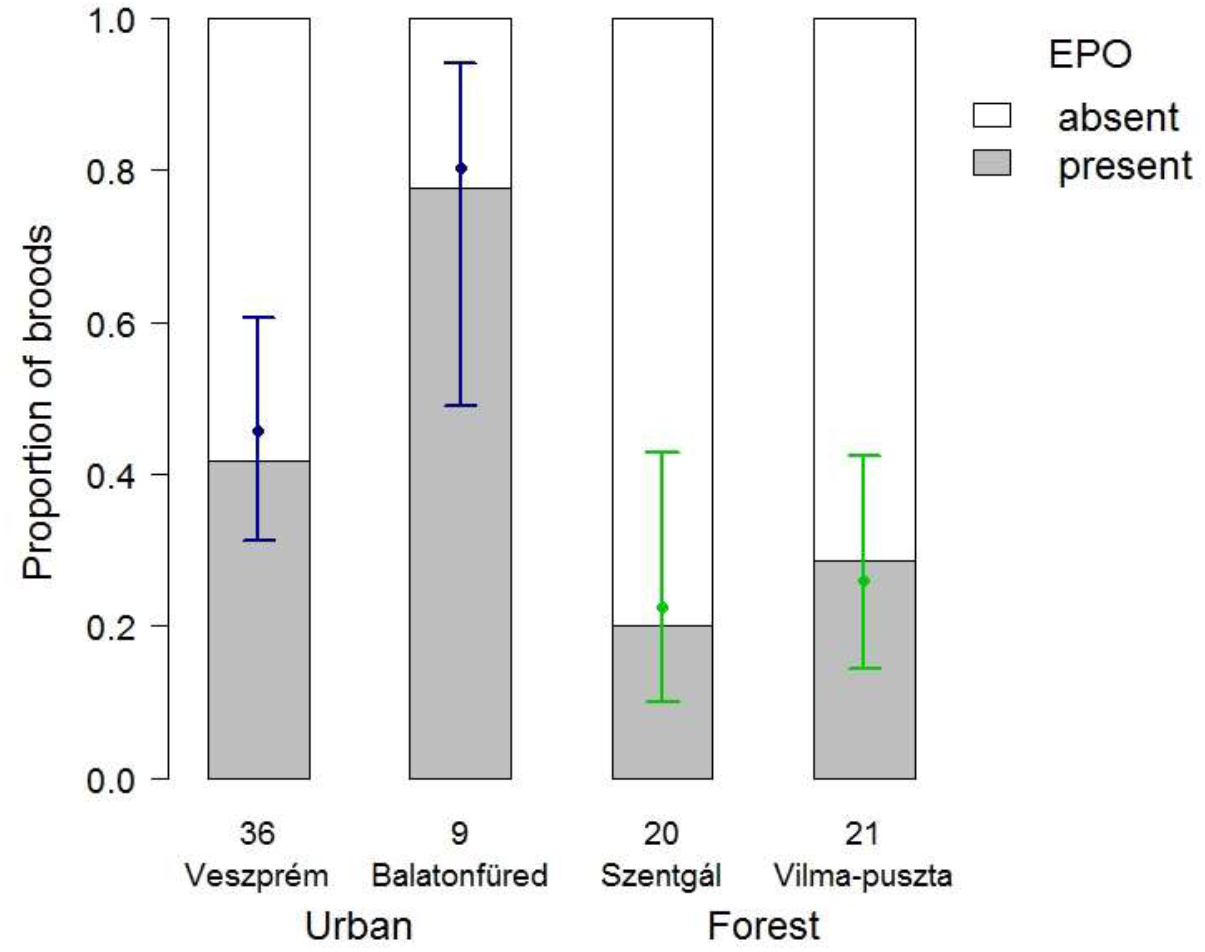
Occurrence of extra-pair offspring (EPO) in great tit broods at urban and forest study sites. Numbers below the bars refer to the number of genotyped broods in each site. Dots and whiskers show means and 87% confidence intervals, respectively, both calculated from the GEE model with study sites and years as predictors. Non-overlapping 87% CIs indicate statistically significant difference (i.e. that a 95% CI of the difference excludes zero).

**Table 1.**
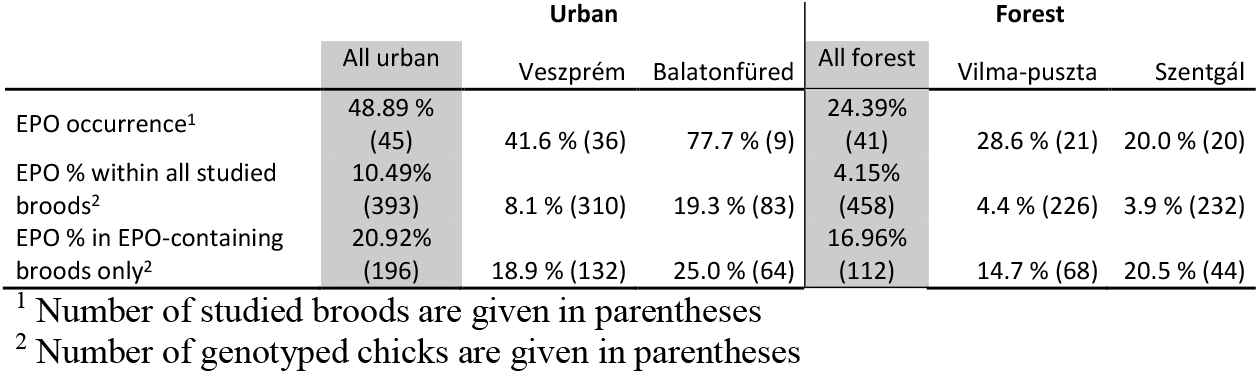
Occurrence of extra-pair offspring (EPO), i.e. proportion of studied broods that contained at least one EPO, and the percentage of EPO among all nestlings in the two urban and two forest study sites.

The number and proportion of EPO was slightly higher in urban broods than in forests when we considered all broods (Table 1), but when we considered only the broods with at least one EPO there was no difference between urban and forest habitats in the number of EPO per brood (Table 1, Figure S1, Table S2, proportional difference between urban and forest broods from linear contrast = 1.175, 95 % CI: 0.667 – 2.067, p = 0.58) and the proportion of EPO per brood (Table 1, Figure S2, Table S2, odds ratio = 1.65, 95 % CI: 0.793 – 3.43 from urban-forest linear contrast, p = 0.18).

We detected two events of intraspecific brood parasitism in 2014 with 2 out of 9 (Veszprém, urban site) and 3 out of 13 nestlings (Szentgál, forest site) being mismatched with their social mother’s genotype on at least two loci. We also found one case of interspecific brood parasitism in one urban nestbox in Veszprém in 2013, where a blue tit (*Cyanistes caeruleus*) nestling was reared successfully in a great tit brood.

## Discussion

We found that urban broods of great tits contained at least one EPO significantly more often than forest broods, and this pattern was consistent across all three years. This result corroborates findings on other species: EPF tended to be higher in more urbanized areas in Canada geese (*Branta canadensis*) (Moore *et al*. 2012) and an unusually high rate of EPF was registered in urban Cooper’s hawks (*Accipiter cooperii*) (Rosenfield *et al*. 2015). However, there was no habitat difference in EPF rate of mountain chickadees (*Poecile gambeli*) (Bonderud *et al*. 2018). On the other hand, in broods where EPO occurred we did not find difference in the number and proportion of EPO between urban and forest habitats. Thus our results suggest that urbanization is associated with increased occurrence of EPF in great tits, although those females that engage in EPF produce similar numbers of EPO in both habitat types.

Several factors can explain the higher occurrence of EPO in urban populations. First, urban sites may offer more opportunities in space and time, because breeding sites are often limited and more fragmented in cities, which can cause higher breeding density in the suitable habitat patches (Crooks *et al*. 2004; Moller *et al*. 2012), and higher breeding density can increase EPF frequency (Charmantier & Perret 2004; Mayer & Pasinelli 2013). Moreover, urban individuals often have a prolonged diurnal activity due to artificial lighting at night (Dominoni *et al*. 2013), offering more time for females to search for extra-pair males during early dawn (Double & Cockburn 2000; Kempenaers *et al*. 2010; Halfwerk *et al*. 2011). Furthermore, night lighting enables males to start singing earlier and such males are more successful in siring EPO (Silva, Valcu & Kempenaers 2015).

Further, food availability and its seasonal distribution may also contribute to the differences in extra-pair mating behaviour between urban and forest populations. Caterpillars, the main food of great tits during the breeding season, are scarce in cities (Biard *et al*. 2017; Seress *et al*. 2018), so urban males might have to spend more time foraging at the expense of guarding their females during their fertile period, increasing opportunities for EPF in cities. However, previous experimental studies found that food scarcity can either increase (Hoi-leitner *et al*. 1999) or decrease EPF rates in various species (Václav, Hoi & Blomqvist 2003; Kaiser *et al*. 2015). Furthermore, urban habitats lack the massive spring peak of caterpillar abundance that is typical in forests (Seress *et al*. 2018), which might lead to less synchronized breeding in urban populations, further facilitating EPF (Van Dongen & Mulder 2009; García-Navas *et al*. 2015), although an opposite effect of breeding asynchrony is also possible (Stutchbury & Morton 1995; Neudorf 2004).

Also, birds might gain more benefits from EPF in urban habitats than in forests. Urban birds can suffer from a higher risk of inbreeding (Vangestel *et al*. 2011; Rodriguez-Martínez *et al*. 2014) and reduced reproductive success per breeding attempt (Bailly *et al*. 2016; Biard *et al*. 2017; Seress *et al*. 2018), so any benefit gained from fertility insurance and good genes might have more value in urban areas (Reding 2014). Furthermore, urban females may be more motivated to pursue extra-pair partners because they might be more likely to perceive their social mate’s quality as low. Individual quality and/or signals like song and plumage ornaments in birds are often relatively poorly developed in urban habitats, including the great tits’ yellow plumage coloration (Halfwerk *et al*. 2011; Biard *et al*. 2017), black breast stripe (Senar *et al*. 2014) and song characteristics (Slabbekoorn & Peet 2003). Urban great tit males that sing higher-frequency songs to overcome the noise-induced communication breakdown are cuckolded more often (Halfwerk *et al*. 2011), although another study found no difference in EPF in house sparrows (*Passer domesticus*) between noisy and quiet breeding places (Schroeder *et al*. 2012).

Alternatively, the relationship between urbanization and EPO occurrence might be non-causal. For example, individuals with different behavioural types can differ in their habitat choice (Holtmann *et al*. 2017; Sprau & Dingemanse 2017) and the propensity to engage in EPF can vary between females as part of their personality (Forstmeier 2007; Forstmeier *et al*. 2014; Wolak *et al*. 2018). It is thus possible that a tendency for promiscuity is associated with the behavioural traits that facilitate settlement in cities, such as innovative problem solving and exploratory behaviour (Bókony *et al*. 2017).

Taken together, there are many conceivable mechanisms by which habitat urbanization may influence the frequency of EPF. Understanding how these mechanisms shape avian reproductive behaviours, and why their effects differ between species, will further expand our knowledge on urban behavioural ecology and sexual selection. Furthermore, because EPF can decrease inbreeding and increase genetic diversity, uncovering the genetic mating systems of urban animals and identifying the drivers of urban cuckoldry may aid the conservation of fragmented populations in our urbanizing world.

## Acknowledgements

We thank T. Hammer and S. Papp for helping with field work. The project was supported by the European Union, with the co-funding of the European Social Fund (TÁMOP-4.2.2.A-11/1/KONV-2012-0064) and by the Hungarian Scientific Research Fund (NKFIH K84132, K112838). IP was supported by the ÚNKP-17-3 New National Excellence Program of the Ministry of Human Capacities, Hungary. VB was supported by the János Bolyai Scholarship of the Hungarian Academy of Sciences. GS was supported by an NKFIH postdoctoral grant (PD 120998) during the preparation of the manuscript.

## Additional information

The authors declare that they have no conflict of interest.

## Author Contributions

IP, VB, KS and AL designed the study. IP, VB, BP, GS, EV and AL collected data in the field. IP and KS did the molecular work and read the genotypes. KS made the parentage analyses. IP, VB and AL did the statistical analyses. All authors wrote the manuscript.

## Supplementary Material

### Field methods

Urban study sites are located in the inner parts of the city of Veszprém (47°05’17.29”N, 17°54’29.66”E) and Balatonfüred (46°57’30.82”N, 17°53’34.47”E), mostly in public parks, university campuses and a cemetery, where vegetation contains both native and introduced plant species. The nest-box plot in Balatonfüred was set up at the end of 2012, so we have data from 2 breeding seasons for this site. Forest study sites are located in deciduous woodlands near Szentgál (47°06’39.75”N, 17°41’17.94”E; characterized mainly by beech *Fagus sylvatica* and hornbeam *Carpinus betulus*) and Vilma-puszta (47°05’06.7”N, 17°51’51.4”E; characterized mainly by downy oak *Quercus cerris* and South European flowering ash *Fraxinus ornus*).

When we monitored the broods, we captured parent birds using a nest-box trap 6-15 days after their first nestling had hatched. We determined parents’ sex based on their plumage characteristics, and ringed each bird with a unique combination of a numbered metal ring and three plastic colour rings. We collected a small amount of blood (ca. 25 μl) from the brachial vein into 500 μl Queen’s lysis solution or 96 % ethanol. When the nestlings reached the age of 14-16 days (mean ± SE: 15.19 ± 0.05 days) we ringed them and took blood samples using the same methods as with their parents. In 2013 and 2014 we also collected the eggs that did not hatch five days after the first nestling had hatched, as well as tissue samples from the nestlings that were found dead before blood sampling, and stored them in 96 % ethanol. All procedures were in accordance with the guidelines for animal care outlined by ASAB/ABS (www.asab.org) and Hungarian laws, licensed by the Government Office of Veszprém County, Nature Conservation Division (former Middle Transdanubian Inspectorate for Environmental Protection, Natural Protection and Water Management; permission number: 31559/2011).

### Genotyping

The samples were stored at 4°C until their laboratory use. For genotyping, we used only the first annual brood of selected pairs: in each study site, a clutch was regarded as first brood if it was initiated before the date of the first egg laid in the earliest second clutch at that site by an individually identifiable (i.e. colour-ringed) female that had a successful first breeding (i.e. fledged at least one young) in that year. We extracted DNA using silica membrane isolation kits (GeneJET Genomic DNA Purification Kit, ThermoFisher Scientific). Forward primers were labelled with fluorescent dyes (Fam-6, NED, PET, or HEX) on the 5’ end; reverse primers contained a GTTT pigtail sequence on their 5’ end. PCR reactions were performed in 20 μl volumes, containing 10-30 ng of total genomic DNA template, 1 U of DreamTaq polymerase (Fermentas), 1 × DreamTaq PCR buffer (Fermentas), 1.5 mM MgCl_2_, 10 pmol dNTPs (Fermentas) and 10 pmol of the respective primer(s). PCR profiles were the following for all loci: initial denaturation at 95°C for 2 minutes, followed by 39 cycles of 95 °C for 30 sec, 57 °C for 45 sec and 72 °C for 45 sec, concluded by a final extension step at 72°C for 7 min. In a subset of samples (n=170 taken in 2014), direct PCR method was used to reduce the cost of laboratory work. Direct PCR process was done without DNA extraction, using the following PCR profile: initial denaturation at 98°C for 3 minutes, followed by 40 cycles of 98 °C for 6 sec, 56 °C for 15 sec, 72 °C for 20 sec. Direct PCR reactions were made in 20 μl volumes containing 1 μl of diluted blood template, 10 μl of Phire Tissue Direct PCR Master Mix (Thermo Fisher Scientific, USA) and 10 pmol of the respective primer(s). Fluorescent PCR products were scanned by capillary electrophoresis on an Abi 3130 Genetic Analyser (Thermo Fisher Scientific); alleles were identified and scored with PEAKSCANNER software (Thermo Fisher Scientific) by two independent, experienced researchers who were blind to the identity of individuals. We successfully genotyped all blood samples (n = 972 in total, 159 adults and 819 nestlings) and all tissue samples from dead nestlings (n = 17). We found 16 embryos in the 46 unhatched eggs, and 15 (93.8 %) embryos were successfully genotyped (see more on the unhatched eggs below). There were nine parents with reconstructed genotypes (Jones *et al*. 2010) and 14 parents with more than one of their broods genotyped in different years (n = 21 broods).

The 5 microsatellite markers were highly variable (see Table S1 for diversity indices). Both expected and observed heterozygosities for the 5 loci averaged 0.88 (SD = 0.03). The probability of identity when siblings were present was 2.99 × 10^−3^ for the 5 loci combined and 1.42 × 10^−4^ for the 8 loci combined. Using MICROCHECKER 2.2.2 (Van Oosterhout *et al*. 2004) we did not find evidence for large-allele dropout and genotyping errors due to stutter bands at any of the 8 loci; null alleles may have been present at one locus (PmaGAn27). Using GENPOP 4.0 (Rousset 2008) we detected no departure from Hardy-Weinberg equilibrium, but significant linkage disequilibrium for 3 pairs of loci (PmaTGAn59 with PmaTAGAn89, PmaTGAn33 with PmaGAn27 and PmaTGAn54; p < 0.001). To further validate our 5-loci marker set, we conducted parentage analysis with CERVUS 3.0 (Kalinowski, Taper & Marshall 2007) using the data of 82 candidate fathers and 58 offspring that had no mismatch with their social father’s genotype, and providing the mothers’ genotype. For each offspring tested, its social father received positive LOD score (i.e. the sum of the log-likelihood ratios at each locus), meaning that this male was more likely to be the genetic father than the other candidate fathers were. When more than one candidate fathers had positive LOD scores (n= 26 offspring), the social father always ranked first, i.e. had the highest LOD score. Thus, our 5-loci marker set proved reliable and efficient for discriminating between within-pair offspring (WPO) and extra-pair offspring (EPO).

The number of paternally mismatched loci per offspring ranged between 2 and 7. We could not identify the genetic father of the majority (47 out of 60) of EPO because we did not have DNA samples from all males in each population. Altogether 159 parents (80 males and 79 females) and 851 offspring (n=819 live chicks, n=17 dead chicks, n=15 embryos in unhatched eggs) were genotyped in 86 families. There were seven males and seven females that each had two of their broods in different years genotyped (n = 21 broods). Out of the 86 broods, 50 were complete broods where all offspring (i.e. all the laid eggs) were genotyped either because every egg hatched and reached the age of blood sampling or because every unhatched egg and/or dead nestling was sampled. For 13 additional broods, we found no embryos in the collected unhatched eggs (n = 30 eggs), so we assumed that these eggs were infertile. The remaining 23 broods were incomplete because some unhatched eggs (n = 12 broods) or dead nestlings (n = 8 broods) or both (n = 3 broods) disappeared from the nest before we could sample them. From the 17 dead nestlings that we genotyped, two were identified as EPO, and there were additional EPO in the same broods among the alive EPO. None of the genotyped 15 embryos were identified as EPO. We found EPO in 62% of the incomplete urban broods (8 / 13 broods) and 44% of the complete urban broods (14 / 32 broods). In forest broods, we found EPO in 20% of incomplete broods (2 / 10 broods) and 26% of the complete broods (8 / 31 broods). The proportion of incomplete broods did not differ between urban (13 / 45) and forest (10 / 41) broods (Fisher’s exact test: odds ratio: 1.26, p = 0.808), and the frequency of EPO in incomplete broods was not different from the frequency of EPO in complete broods (Fisher’s exact test, urban broods: odds ratio: 2.02, p = 0.337, forest broods: odds ratio: 0.72, p > 0.99). Thus, incomplete broods are unlikely to introduce bias in our data, and considering the missing unsampled offspring in incomplete broods as WPO makes our estimates of EPO occurrence conservative, so we analysed complete and incomplete broods together.

### Statistical analyses

We tested the difference in EPO frequencies between urban and forest broods by Generalised Estimation Equations (GEE) models. We allowed for the non-independence of those few broods that had at least one parent in common using the ‘exchangeable correlation’ association structure (Zuur *et al*. 2009). We investigated three response variables. (1) For testing the difference in the proportion of broods that contained EPO, the response was a binary variable of EPO presence in a brood (EPO present / absent), and a GEE model was constructed with binomial error structure and logit link function (n = 86 broods). As clutch size varies between urban and non-urban birds (Seress *et al*. 2018), we compared the quantity of EPO between urban and forest broods in two ways, analyzing (2) the number of EPO in a brood using a GEE model with Poisson distribution and log link function, and (3) the proportion of EPO within the brood, calculated as EPO / (EPO + WPO), using a GEE model with binomial error structure and logit link function (n = 32 broods, i.e. only those containing at least one EPO). We included the main effects of study site (4 sites) and year (3 years) in all models (note that testing the site × year interaction was not feasible because we have no data for Balatonfüred from 2012). To statistically compare the two habitat types, we calculated a linear contrast from each of the three GEE model’s estimates (i.e. the difference between the two urban sites and the two forest sites, on log scale). Each linear contrast was back-transformed from the log-scale to provide the odds ratio (i.e. proportional difference of the odds of having at least one EPO, or a chick being EPO, between urban and forest broods) for EPO presence and EPO proportion and the incidence rate ratio (i.e. proportional difference of EPO frequency between urban and forest broods) for EPO number. All analyses were implemented in the R 3.1.1 software environment (R Core Team 2014), using packages “geepack” (Højsgaard, Halekoh & Yan 2006), “emmeans” (Lenth 2018) and “multcomp” (Hothorn, Bretz & Westfall 2008).

## Supplmentary Figures

**Figure S1.**
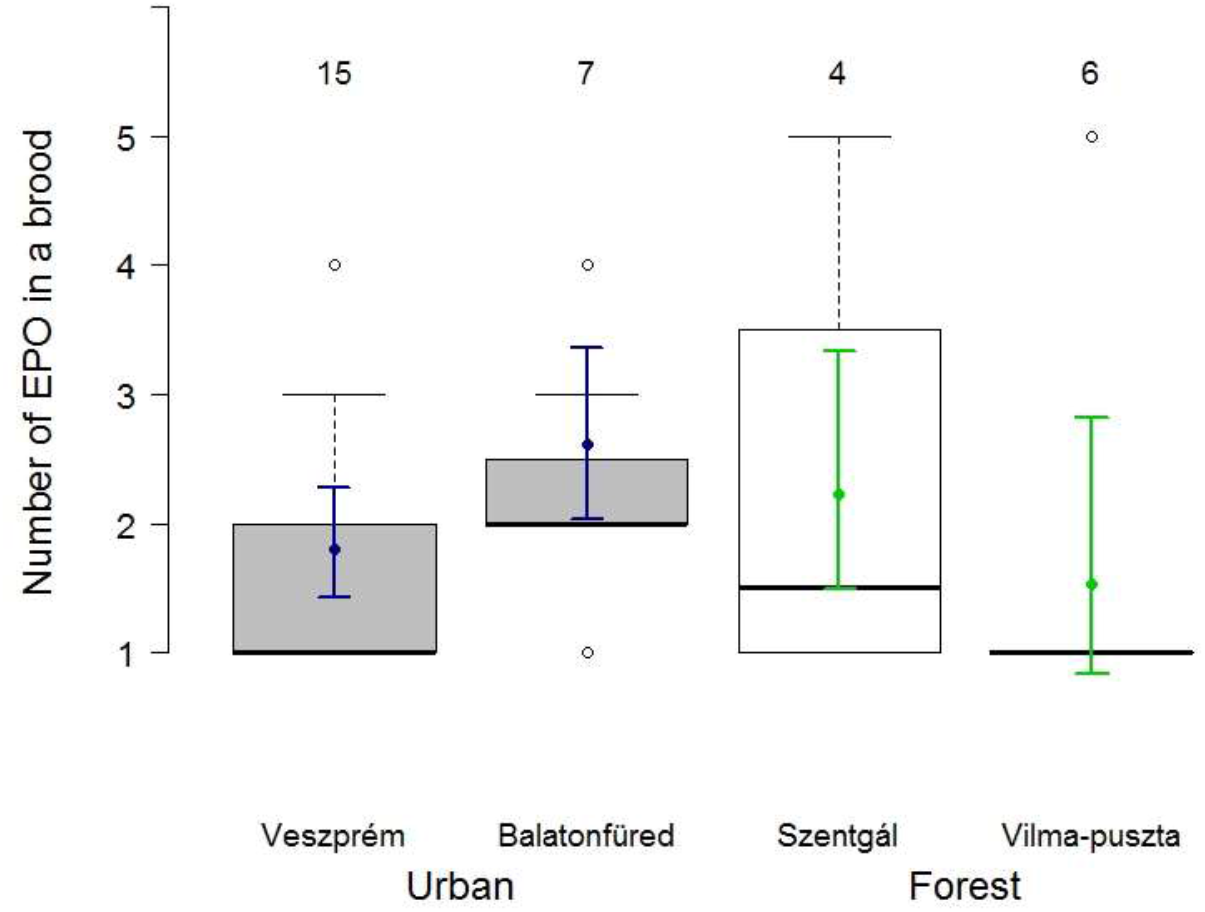
Number of extra-pair offspring (EPO) in EPO-containing broods in our study sites. Numbers above the boxes refer to the number of genotyped broods in each site. In each boxplot, the thick middle line and the box shows the median and the interquartile range, respectively; dashed whiskers extend to the most extreme data points within 1.5 × interquartile range from the box, and empty dots represent more extreme data points. Coloured filled dots and solid whiskers show the mean and its 87% confidence interval, respectively, both calculated from the GEE model with study sites and years as predictors. Non-overlapping 87% CIs indicate statistically significant difference (i.e. that a 95% CI of the difference excludes zero). The number of EPO was one in n=9 urban and n=7 forest broods, two in n=9 urban and n=1 forest broods. There were n=2 urban broods with three EPO, n=2 additional urban broods with four EPO, and n=2 forest broods with five EPO.

**Figure S2.**
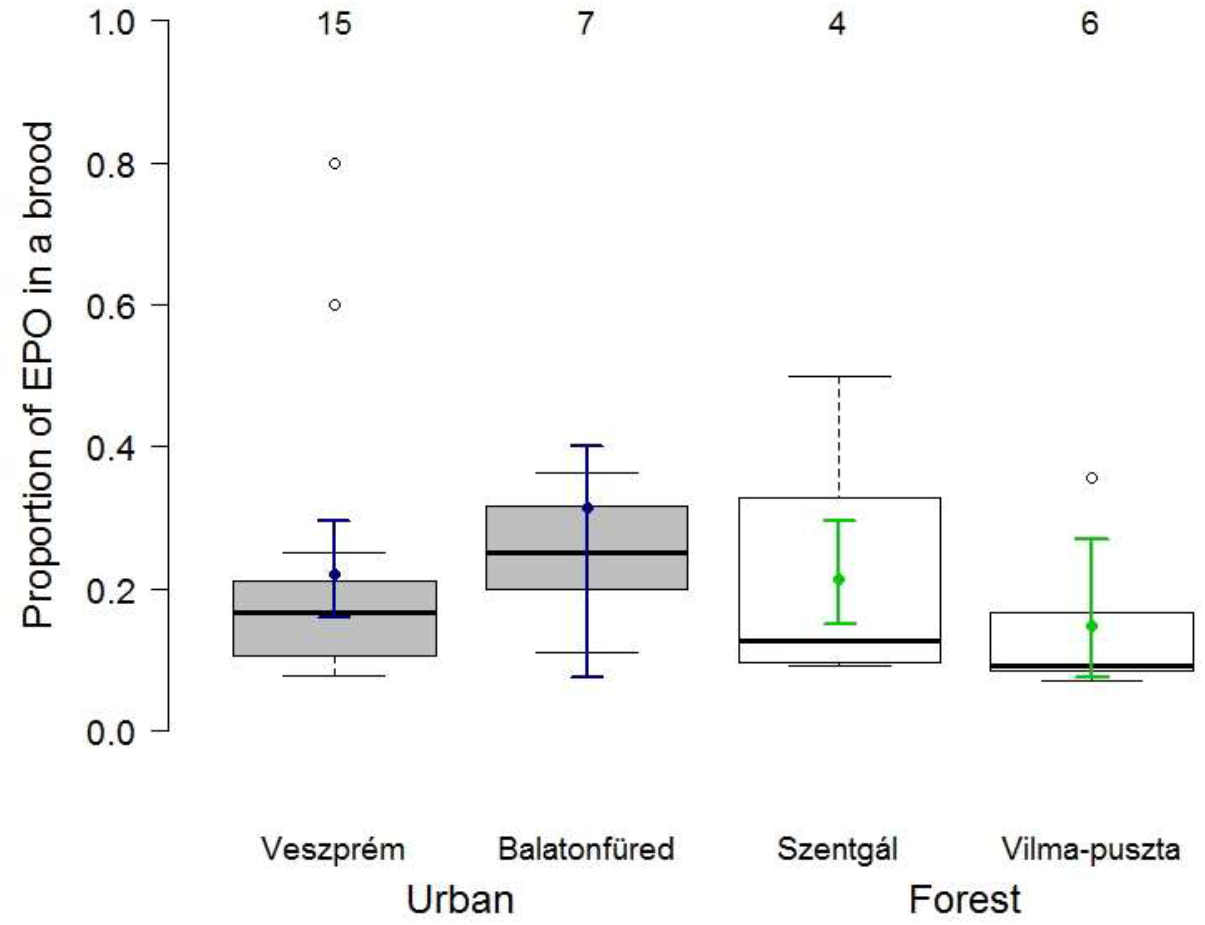
Proportion of extra-pair offspring (EPO) in EPO-containing broods in our study sites. Numbers above the boxes refer to the number of genotyped broods in each site. In each boxplot, the thick middle line and the box shows the median and the interquartile range, respectively; dashed whiskers extend to the most extreme data points within 1.5 × interquartile range from the box, and empty dots represent more extreme data points. Coloured filled dots and solid whiskers show the mean and its 87% confidence interval, respectively, both calculated from the GEE model with study sites and years as predictors. Non-overlapping 87% CIs indicate statistically significant difference (i.e. that a 95% CI of the difference excludes zero). The proportion of EPO where at least one EPO occurred ranged from 0.08 to 0.80 in urban broods (median: 0.2) and between 0.07 and 0.50 in forest broods (median: 0.095).

**Table S1.** Observed allele diversity, probability of identity (PI), probability of exclusion with both parents known (PE2) and with only one parent known (PE1), and GenBank accession number of the microsatellite loci used in the study

| Locus | No. of alleles | PI | PE2 | PE1 | GenBank |
| --- | --- | --- | --- | --- | --- |
| PmaGAn27 | 22 | 0.011 | 0.846 | 0.732 | AY260532 |
| PmaTAGAn89 | 14 | 0.035 | 0.719 | 0.558 | HQ263126 |
| PmaTGAn33 | 16 | 0.023 | 0.773 | 0.630 | AY260539 |
| PmaTGAn54 | 32 | 0.031 | 0.733 | 0.576 | HQ263130 |
| PmaTGAn59 | 20 | 0.016 | 0.814 | 0.686 | HQ263131 |
| **5 loci combined** | **104** | **$4.26 \times 10^{-9}$** | **>0.999** | **0.994** |  |
| PmaCAn1* | 11 | 0.036 | 0.715 | 0.554 | AY260530 |
| PmaTAGAn73* | 9 | 0.054 | 0.653 | 0.480 | HQ263122 |
| PmaTAGAn78* | 17 | 0.014 | 0.823 | 0.700 | HQ263123 |
| **8 loci combined\*** | **141** | **$1.43 \times 10^{-12}$** | **>0.999** | **0.999** |  |

The data are based on 159 adults for the first five loci and 23 adults for the three loci marked with an asterisk.

**Table S2.** Results of Generalised Estimation Equations (GEE) models for the occurrence, number and proportion of EPO in great tit broods. Study site (4 sites) and year (3 years) were the predictor variables in the models. Exponentially back-transformed parameter estimates (**e^b^**) for the four sites provide the odds of having at least one EPO in a brood, the frequency of EPO in broods that contained EPO, and the odds of each offspring being EPO in broods that contained EPO, in the year 2014. For years 2012 and 2013, **e^b^** provides the difference from 2014, i.e. the odds ratio of having at least one EPO, the proportional difference of the frequency of EPO, and the odds ratio of each offspring being EPO. Parameter estimates (**b**) with their standard errors (**SE**) from the GEE models are also presented, and n is the number of genotyped broods in the analyses.

| Model parameters | EPO occurrence (n=86) |  |  |  | EPO number (n=32) |  |  |  | EPO proportion (n=32) |  |  |  |
| --- | --- | --- | --- | --- | --- | --- | --- | --- | --- | --- | --- | --- |
| | b ± SE | $e^b$ | $\chi^2$ | p | b ± SE | $e^b$ | $\chi^2$ | p | b ± SE | $e^b$ | $\chi^2$ | p |
| **site: Veszprém** | 0.301 ± 0.466 | 0.575 | 0.42 | 0.519 | 0.541 ± 0.140 | 1.720 | 14.89 | < 0.001 | -1.589 ± 0.187 | 0.170 | 72.33 | < 0.001 |
| **site: Balatonfüred** | 1.877 ± 0.916 | 0.867 | 4.20 | 0.040 | 0.911 ± 0.152 | 2.490 | 35.84 | < 0.001 | -1.110 ± 0.191 | 0.248 | 33.74 | < 0.001 |
| **site: Szentgál** | -1.764 ± 0.608 | 0.318 | 1.58 | 0.209 | 0.753 ± 0.259 | 2.120 | 8.41 | 0.473 | -1.626 ± 0.301 | 0.164 | 29.25 | < 0.001 |
| **site: Vilma-puszta** | -0.564 ± 0.586 | 0.362 | 0.93 | 0.336 | 0.377 ± 0.465 | 1.460 | 0.66 | 0.715 | -2.073 ± 0.582 | 0.112 | 13.12 | < 0.001 |
| **year: 2012** | -0.291 ± 0.715 | 0.428 | 0.17 | 0.684 | 0.357 ± 0.326 | 1.430 | 1.20 | 0.273 | 0.948 ± 0.546 | 0.721 | 3.02 | 0.082 |
| **year: 2013** | -1.142 ± 0.460 | 0.242 | 6.18 | 0.013 | -0.209 ± 0.186 | 0.811 | 1.25 | 0.263 | 0.033 ± 0.272 | 0.508 | 0.01 | 0.905 |

